# On Motivating the Choice of Link Function in Binomial Regression Through A Latent Abundance Process

**DOI:** 10.64898/2026.10.16.759589

**Authors:** Bert van der Veen, Francis K.C. Hui

## Abstract

Ecologists commonly apply binomial regression to study binary data. Often, the link function in binomial regression is not purposefully chosen, but instead selected on the grounds of interpretation or computation efficiency. However, if the binary response is the result of a latent abundance process, then the link function implies assumptions for this underlying process. The most well-known example of this is the complementary log-log (cloglog) link, which has direct connections with the probability of a positive count under a latent Poisson distribution. On the other hand, counts in ecology are often overdispersed, meaning the assumption of an underlying Poisson abundance process is unrealistic.

In this article, we provide a framework for consideration of the link choice in binomial regression, in terms of the latent abundance process that binary responses can represent. Aside from the Poisson case mentioned above, we show that the cloglog link is the right choice if the latent abundance process represents the biomass of individuals, or if the latent abundance process comes from an overdispersed count distribution with a linear mean-variance relationship. For the case where the latent count follows a negative binomial distribution with a quadratic mean-variance relationship, we derive a new generalized complementary log-log (gcloglog) link function, which includes the case of the cloglog link as well as that of the logit link, where the latter implies assuming moderate overdispersion in the form of a geometric distribution. Two simulation studies demonstrate that cloglog and logit link binomial regression can exhibit bias and undercoverage when the latent count is more overdispersed than assumed, while gcloglog link models overcome this issue but in practice require replication in the binomial response for reliable estimation. Two real ecological datasets further illustrate how to use the gcloglog link binomial regression in practice including as a means of sensitivity analysis. We encourage ecologists to give more thought to the choice of link function in binomial regression, based on considering the latent abundance process and its properties.

## 1 Introduction

Binomial regression is the tool of choice when it comes to analyzing binary or bounded responses in ecology, with a myriad of applications ranging from modeling the prevalence of species in (joint) species distribution modeling (Elith and Leathwick, 2009; Wilkinson et al., 2019), model-based ordination and clustering (Dunstan et al., 2013; Warton et al., 2015; Thorson et al., 2016; van der Veen et al., 2023), successful germination of seeds out of a total (Crowder, 1978; Mackenzie et al., 2016), and habitat selection (Fieberg et al., 2021), among many others.

While considerable thought is often given to the decision around what terms are to be included in the linear predictor of a binomial regression (Tredennick et al., 2021), an equally important but often overlooked decision is the choice of link function connecting the mean of the binomial distribution to the linear predictor. For the vast majority of applications in ecology, the canonical logit link is used, while in Bayesian statistics the probit link has long been favored due to its amenability with a data augmentation scheme that facilitates Markov-Chain Monte-Carlo sampling (Albert and Chib, 1993), although recent years have also witnessed an increase in the use of the logit link due to the Polya-gamma augmentation scheme (Polson et al., 2013). A third, less commonly used link function is the complementary log-log link (abbreviated in this article to cloglog link), which is grounded in the truncation of a latent Poisson count; see the next paragraph for further discussion. Other link functions such as the log-link (giving rise to relative risk regression sporadically seen in epidemiology, Marschner 2015) and the family of link functions by Aranda-Ordaz (1981); Jiang et al. (2013), also exist, but are scarcely used in ecology due to computational constraints and/or lack of readily available software that can accommodate flexible linear predictor constructions.

The appropriate choice of link function in a binomial regression depends on the data type and (latent) data generating process. In ecology, an argument can be made for choosing it by considering the nature of the process generating the binary response. For instance, the cloglog link is often used when the latent abundance process is assumed to follow a Poisson distribution, with the binary responses arising from truncating this count such that a zero represents a zero count and a one represents a positive count (Yee and Dirnböck, 2009). Put another way, the cloglog link has a direct connection to the probability of counts under a Poisson distribution, and its usage in a binomial regression suggests an assumption of equivariance for the latent abundance process. This motivation of the cloglog link is well known in the statistical ecology literature (e.g., Yee and Dirnböck 2009; Bowler et al. 2019; Morera-Pujol et al. 2023), although it is sparsely utilized in practice by ecologists. This is perhaps because the connection with a Poisson count process is of limited value given that, in truth, many count processes in ecology are overdispersed and more suitably modeled using a negative binomial distribution (Warton, 2005; Stoklosa et al., 2022), among possible count distribution choices.

When modeling count data, fitting a Poisson regression to overdispersed counts results in biased parameter estimates and typically anticonservative inference (e.g., inflated Type-I errors, Harrison 2014). By extension then, modeling binary data as if coming from a latent Poisson process, when in truth it derives from an overdispersed count mechanism, is likely to provide inaccurate results if the aim is to recover the underlying covariate effects. With this in mind, there is a window of opportunity for improved motivation and dialogue in the broader ecological community around link choice in binomial regressions, with explicit connection to the latent abundance and overdispersed count processes.

In this article, we encourage ecologists to give more thought to the choice of link function in binomial regression, based on deeper consideration of the assumptions around the latent abundance process. We illustrate this by examining the consequences when analyzing binary data with a cloglog link function, when the latent abundance process is truly Tweedie distributed (motivated by latent abundance representing biomass of individuals, Foster and Bravington 2013), or negative binomially distributed (motivated by the latent abundance representing overdispersed counts of individuals, Stoklosa et al. 2022). In the former, we show that the cloglog link arises as the appropriate choice for binary data from latent biomass, although the parameter estimates are not directly interpretable in terms of mean biomass. In the negative binomial case, we further distinguish between a linear and a quadratic mean-variance relationship (Hilbe, 2011), where in the latter we derive a so-called “generalized complementary log-log” or gcloglog link function which, aside from being a more flexible link involving an additional shape parameter, reveals a lesser known result in ecology for the logit link function and what it potentially implies about the latent abundance process.

Through a series of simulations and two real-world case studies, we demonstrate the consequences of misspecifying the link function in binomial regression. In particular, misspecification of the link function can not only induce severe bias in the parameter estimates, but also result in substantial undercoverage of the associated confidence intervals, when the aim is to recover covariate effects in the latent count process. For the derived gcloglog link function, we provide software in `R` for profiling the additional shape parameter and hence link choice, which can straightforwardly be combined with existing Generalized Linear Model (GLM, Nelder and Wedderburn 1972) and Generalized Linear Mixed effects Model (GLMM, Bolker et al. 2009) software to facilitate adoption by ecologists. Besides data-driven estimation of the shape parameter, the software also serves as a tool for sensitivity analysis to assess the severity of misspecifying the link function especially in smaller sample sizes.

This reflects our broader goal of encouraging greater scrutiny of the link choice in binomial regression, and acknowledging the more flexible gcloglog link function may not always be the best choice for the application of interest.

The remainder of this article is structured as follows. Section 2 reviews binomial regression and draws connections between the binary response and Tweedie and negative binomial distributions for latent abundance. Section 3 introduces the gcloglog link function, quantifies the information lost by truncating an overdispersed count, and presents the accompanying `R`-package. Section 4 presents the numerical studies to explore the practical data requirements of the gcloglog link function, and consequences of fitting logit or cloglog regression when the latent count is overdispersed. Finally, in Section 5 we apply the gcloglog link to two real-world datasets on bats (Cole, 2022) and salamanders (Price et al., 2016). The `gcloglog` package is available at https://github.com/BertvanderVeen/gcloglog.

## 2 Binomial data in terms of a latent abundance process

Let *y*_*i*_ ~ Binom(*N*_*i*_, *p*_*i*_) denote the response at observational unit *i*, which we assume to follow a binomial distribution with trial size *N*_*i*_ and probability of success (or equivalently the mean) *p*_*i*_. The latter is regressed as a function of covariates ***x***_*i*_ as

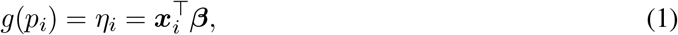

where ***β*** denotes the corresponding regression coefficients. The above binomial regression can be extended in a myriad of ways, e.g., including random effects, smooth terms, and so on, with the developments below applying to these more advanced settings as well. Finally, *g*(*·*) denotes the link function that connects the probability of success to the linear predictor *η*_*i*_. As reviewed in Section 1, the three most commonly used links for binomial regression in ecology are:

- Logit link: *g*(*p*_*i*_) = log*{p*_*i*_*/*(1 *− p*_*i*_)*}*. Aside from being the canonical link function for the binomial distribution, this link offers an interpretation of the covariate effects in terms of oddsratios.
- Probit link: *g*(*p*_*i*_) = Φ^*−*1^(*p*_*i*_), where Φ(*·*) denotes the standard normal cumulative distribution function. The link is often viewed as taking an underlying normal distribution *z*_*i*_ with mean *η*_*i*_ and variance one, and setting *y*_*i*_ = 1 if *z*_*i*_ *≥* 0 and zero otherwise.
- Complementary log-log (cloglog): *g*(*p*_*i*_) = log*{−* log(1 *− p*_*i*_)*}*. Unlike its logit and probit counterparts, the cloglog link is asymmetric around *η*_*i*_ = 0 (equivalently around *p*_*i*_ = 0.5) and approaches zero much slower than one. We discuss this link function in greater detail below.

Let 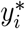 represent a latent abundance process of an ecological phenomenon, such that the binary response introduced above arises from truncating this process and “success” is recorded if the latent abundance is larger than zero. That is, 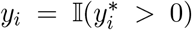 where I(*·*) denotes the indicator function, from which it follows that 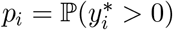.

### 2.1 The complementary log-log link as a latent Tweedie process

Consider first the case where the latent process follows a Tweedie distribution, parametrized by its mean *µ*_*i*_ *>* 0, a dispersion parameter *ϕ >* 0, and a power (or index) parameter *ξ* such that the variance takes the form of a power-law,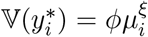. For 1 *< ξ <* 2, the Tweedie distribution provides a flexible way to model biomass, as in this case it can be written in terms of a Poisson sum of gamma random variables and is suited to modeling the total mass of a set of individuals (Foster and Bravington, 2013). Also, note the cases of *ξ* = 0, 1, and 2 correspond to a normal, Poisson, and Gamma distribution, respectively.

Suppose now we regress the mean of the latent Tweedie process using a log link, 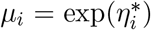, where we use 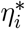 to denote the linear predictor in the latent process. Note the log link is a natural, and by far the most popular, choice when handling non-negative/positive responses. Then it is straightforward to that

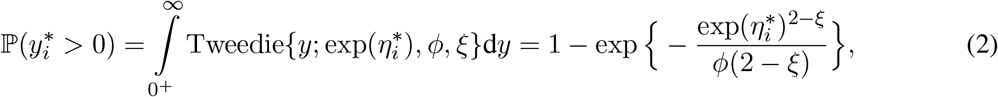

and hence *y*_*i*_ follows a Bernoulli distribution with the probability of success given by the above. Thus if we choose to employ the cloglog link in our binary regression in equation (1), we obtain the relationship

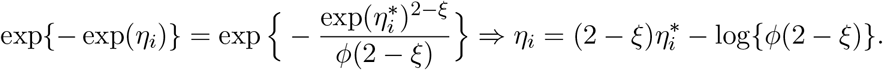

From the above, we see that the cloglog link does not provide immediately interpretable parameters in terms of the mean of the latent Tweedie process, except in the case of the Poisson distribution (as we discuss will shortly). Specifically, if we consider an observed binary response to be the outcome of a latent biomass for multiple individuals, then the cloglog link produces slope effects that are shrunk towards zero relative to the corresponding effects in the latent Tweedie mean by a factor of 0 *<* (2 *− ξ*) *<* 1, while the intercept is shifted from the intercept of the latent Tweedie mean by the constant *−∞ <* log*{ϕ*(2 *− ξ*)*} <* log *ϕ*.

In the case of *ϕ* = *ξ* = 1 the Tweedie distribution reduces to the Poisson distribution, and hence the above reveals equivalence in the linear predictor, 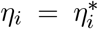. That is, if the latent abundance process 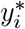 follows exactly a Poisson distribution, then binomial regression with the cloglog link function provides regression coefficients directly interpretable in terms of the Poisson rate (mean) parameter, as is already known in the literature; see Section 1.

### 2.2 Binomial regression with a latent negative binomial process

In practice, count data in ecology are often overdispersed (Warton, 2005; Stoklosa et al., 2022). With this in mind, in this section we assume the latent abundance process follows a negative binomial distribution 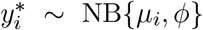 with mean *µ*_*i*_ and overdispersion parameter *ϕ >* 0, where the latter controls the amount of additional variance relative to the Poisson distribution. There are two commonly applied cases of negative binomial regression in ecology: (I) the linear mean-variance case with 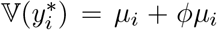, which we refer to here as NBI, and (II) the quadratic mean-variance case where 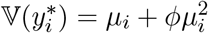, which we refer to here as NBII (Hilbe, 2011).

As in the case of a latent Tweedie process, suppose we regress the mean using a log link, 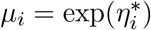 Then if the latent count follows the NBI parametrization, it is straightforward to show that

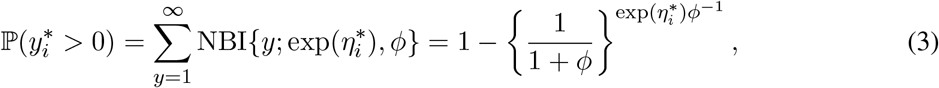

from which it follows that *y*_*i*_ follows a Bernoulli distribution with the probability of success given by the above. Setting the cloglog link equal to the case of equation (3), we obtain

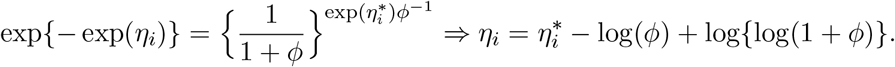

Thus if we consider an observed binary response to be the outcome of a latent count process following a NBI distribution, then the cloglog link is directly able to recover the slope effects of the underlying count mean, but produces an intercept that is biased by a constant value *−∞ <* log(*ϕ*) *−* log*{*log(1 + *ϕ*)*} < ∞*.

Turning to the case of the NBII distribution and the quadratic mean-variance relationship, which is arguably more useful given its greater form of overdispersion (Stoklosa et al., 2022), it is possible to show that

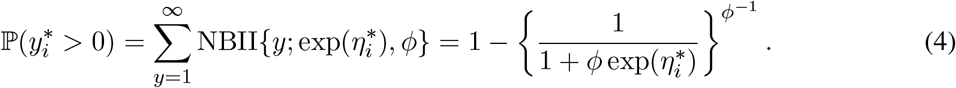

It follows that if we set the mean of a cloglog regression equal to the case of equation (4), we obtain the non-linear form

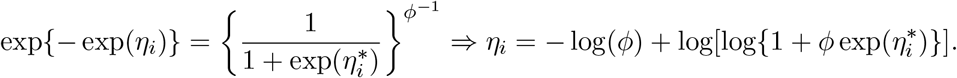

Unlike in the linear mean-variance case, the cloglog link cannot recover the covariate effects of the underlying NBII mean, except in the case where *ϕ →* 0 from which it follows that 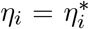 consistent with the discussion at end of Section 2.1. The above result is perhaps not surprising: if we consider an observed binary response to be the outcome of a latent count process following a NBII distribution, then the cloglog link leads to a form of link misspecification since it posits an incorrect (equivariance) shape for the mean-variance relationship of the underlying latent abundance process. We assess the practical consequences of this link misspecification in Section 4.

## 3 The generalized complementary log-log link function

Motivated by the above link misspecification, straightforward algebraic operations can be used to derive the appropriate link function for a binomial regression where the latent overdispersed count follows a NBII distribution. By setting

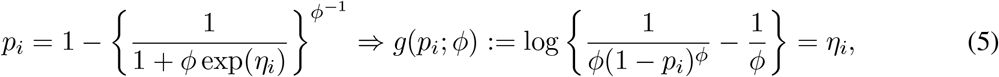

then we refer to *g*(*p*_*i*_; *ϕ*) as the “generalized complementary log-log link” or gcloglog link function, which depends on an additional shape parameter *ϕ >* 0. It is important to acknowledge that the gcloglog link was originally proposed by Aranda-Ordaz (1981), and has been investigated to some degree in the broader statistics literature (e.g., Ntzoufras et al., 2003; Jiang et al., 2013). Unlike these works though, who motivate the gcloglog link largely on the grounds of flexibility, our derivation arrives at the same link function through thinking about the latent abundance process, i.e., the truncation of an overdispersed negative binomial latent count.

In the limit as *ϕ →* 0, the gcloglog link reduces to the cloglog function as expected. A more interesting case arises by setting *ϕ* = 1, such that *g*(*p*; 1) = log*{*1*/*(1 *− p*_*i*_) *−* 1*}* = log*{p*_*i*_*/*(1 *− p*_*i*_)*}*. That is, the popular logit link also arises as a special case of the gcloglog link function, which in turn is a consequence of the NBII distribution simplifying in the case *ϕ* = 1:

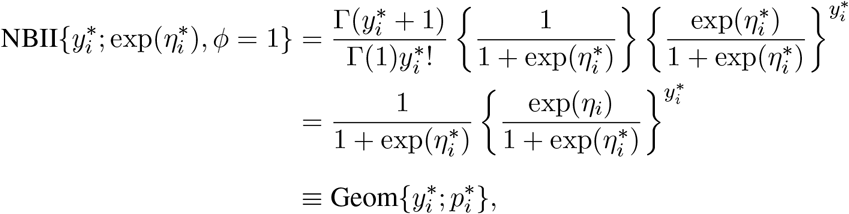

where 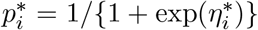. That is, the NBII distribution reduces to the geometric distribution with success probability 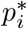. While this equivalence is again well-established in the broader statistical literature, in the context of this article it offers an interesting insight that, just as the usage of a cloglog link function in binomial regression can suggest an assumption of equivariance for the latent abundance process, the usage of a logit link function can suggest an assumption that the latent abundance process is overdispersed with the quadratic form 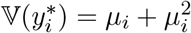.

Figure 1a visualizes the shape of the gcloglog link function for different values of *ϕ*. At *ϕ* = 1 the shape of the link function is the same on both sides of 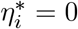 (the logit link), and the rate at which link rises to one and falls to zero is the same either side of this. As *ϕ → ∞* the gcloglog link approaches *p* = 1 much more slowly as a function of the latent count linear predictor. This reflects the more extreme overdispersion with large *ϕ*. Conversely, in the limit as *ϕ →* 0 and we obtain equivariance in the latent count process, the gcloglog link approaches *p* = 0 slowly but rises to *p* = 1 more sharply, reducing to the cloglog link in the limit; see also Agresti (p. 104, 1990).

**Figure 1:**
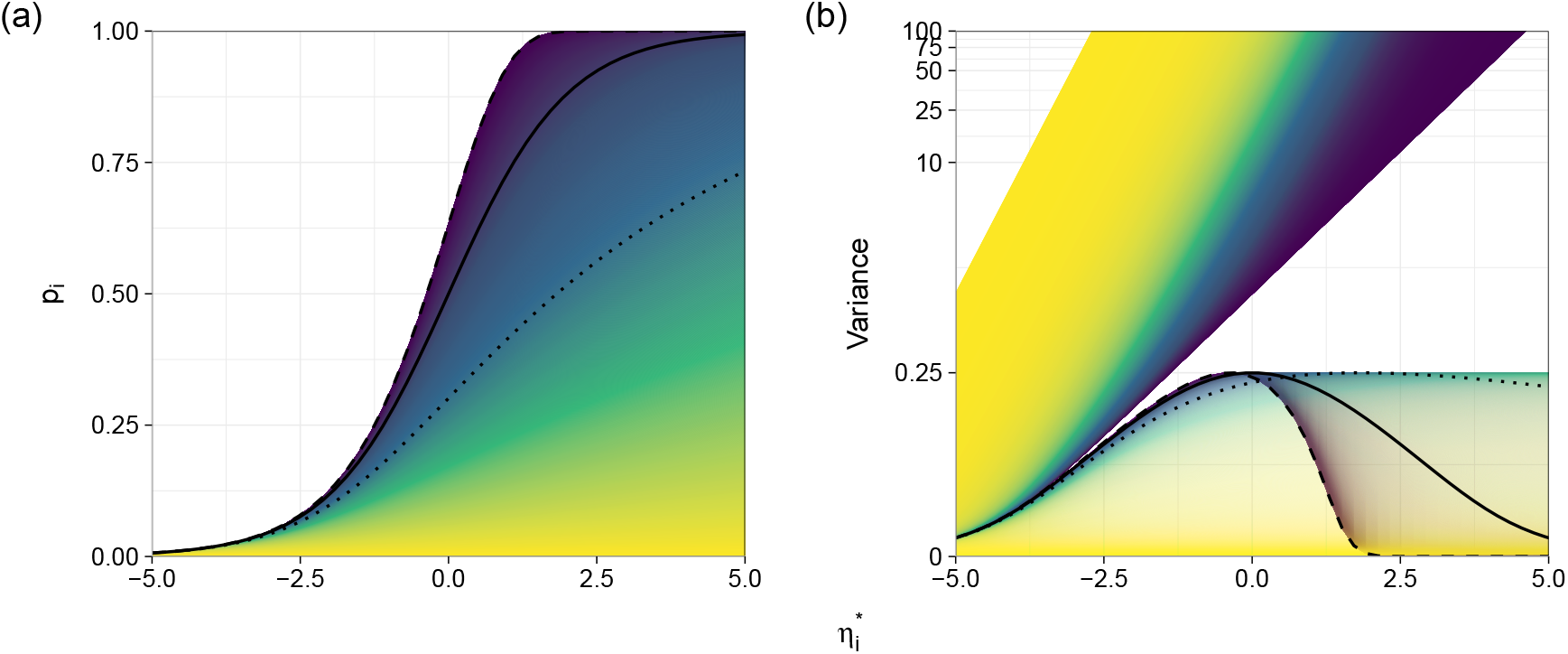
(a) The generalized complementary log-log (gcloglog) function plotted as function of the latent count linear predictor 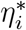, for a range of values for the negative binomial dispersion parameter *ϕ*. A lighter color corresponds to a higher value for *ϕ*, and thus greater overdispersion in the latent count. Superimposed in black are the logit (solid, corresponding to *ϕ* = 1) and complementary log-log (dashed, corresponding to *ϕ* → 0) link functions, while the dotted line represents a case of extreme overdispersion with *ϕ* = 5. (b) Variance of the NBII distribution (top), and of a Bernoulli random variable with gcloglog link (bottom), plotted for the same range of values for *ϕ*. The same three reference lines as in (a) are also included in black. Note transparency has been added for the color of the Bernoulli random variable, to better display overlap of the variance functions for particular combinations of *ϕ* and 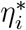.

Yet another way to quantify the effects of how truncating an underlying count process varies with its degree of overdispersion is to examine the relationship in terms of variance. Consider first the case of an underlying Poisson count process with 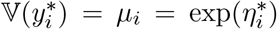. If an observed binary response *y*_*i*_ arises from truncating this count and the cloglog link is used in the resulting binomial regression, then it can be shown that 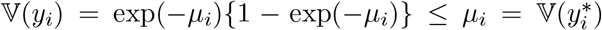 for every 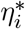. Note the difference between the two variance forms becomes smaller as *µ*_*i*_ *→* 0, meaning that the act of truncation affects the variance of the binary responses corresponding to low mean counts comparably (and unsurprisingly) less than observations corresponding to high mean counts.

The above behavior extends to the case of a latent NBII process. In particular, with a quadratic mean-variance relationship, counts with large *µ*_*i*_ have disproportionately larger variance than counts corresponding to small *µ*_*i*_, a difference that grows with larger *ϕ*. As such, truncation to a binary response *y*_*i*_ de-emphasizes the observations corresponding to the largest variance in the latent count process the most. Figure 1b summarizes this visually, showing how the variance of a NBII distributed random variable behaves as a function of the underlying linear predictor 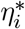, along with the variance of a Bernoulli distributed random variable resulting from the truncating this count and using the gcloglog link function. For *ϕ* = 0, corresponding to the cloglog link, the variance increases relatively slowly until 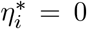, and declines sharply after that, asymptoting to zero as 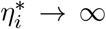 but effectively reaching zero near 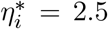. For *ϕ* = 1, corresponding to the logit link, the variance increases to 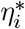 at a similar speed as for the cloglog link, but declines more slowly, effectively reaching zero at 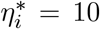. As *ϕ → ∞* the variance function of the latter becomes increasingly heavy right-tailed, i.e., the variance decays much more slowly and the the binomial regression can be thought of as attributing a larger amount of variance for binary responses corresponding to larger *µ*_*i*_, relative to lower *ϕ*.

### 3.1 Information loss

To better understand the practical consequences of how much (or little) binary regression can infer about the covariate effects in the underlying count process, we can compute the so-called “information ratio” between a negative binomial log link regression and the corresponding gcloglog link binomial regression, and examine how this varies with the latent count linear predictor 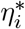 and the degree of overdispersion *ϕ*. Briefly, the information ratio is defined here as the quotient of the Fisher information matrices (expected value of the negative Hessian of the log-likelihood function) between two different models for the same covariate effect. While it is more generally used as a measure of relative statistical efficiency (Lehmann and Casella, 1998), here we compute the information ratio based on the Bernoulli log-likelihood function with gcloglog link, and negative binomial distribution with log link, such that values closer to zero imply more information has been lost due to the truncation process; see Appendix S1 for a formal derivation of this. It is also worth pointing out that the inverse of the Fisher information matrix serves as a standard approach for constructing standard errors in maximum likelihood estimation of GLMs, among other models. As such, the information ratio can also be interpreted as how much narrower Wald-based confidence intervals for slope estimates (say) would be, were the practitioner to have access to the underlying count observations from which the binary response arises. Figure 2 plots the resulting information ratio. Superimposed are contour lines showing values of 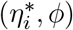 which correspond to the same information loss, along with vertical slides corresponding to the case of *ϕ* = 0 (equivariance, the cloglog link), *ϕ* = 1 (logit link), and *ϕ* = 5 corresponding to more severe overdispersion. Truncating counts with a small mean incurs little loss, as the presence-absence response represents the latent count closely to begin with. In contrast, truncating counts with a large mean loses considerable information, as the presence-absence response is most dissimilar from the latent count. Naturally, there is interplay with the overdispersion parameter, as a large amount of overdispersion worsens the information loss for small mean cases, but lessens it for large mean cases. Put another way, although the interpretation of the slope coefficients from a gcloglog binomial regression is theoretically the same as in a log link NBII regression, the actual estimates and associated uncertainties can, as expected, differ greatly in finite samples given binary data is information poor relative to overdispersed counts.

**Figure 2:**
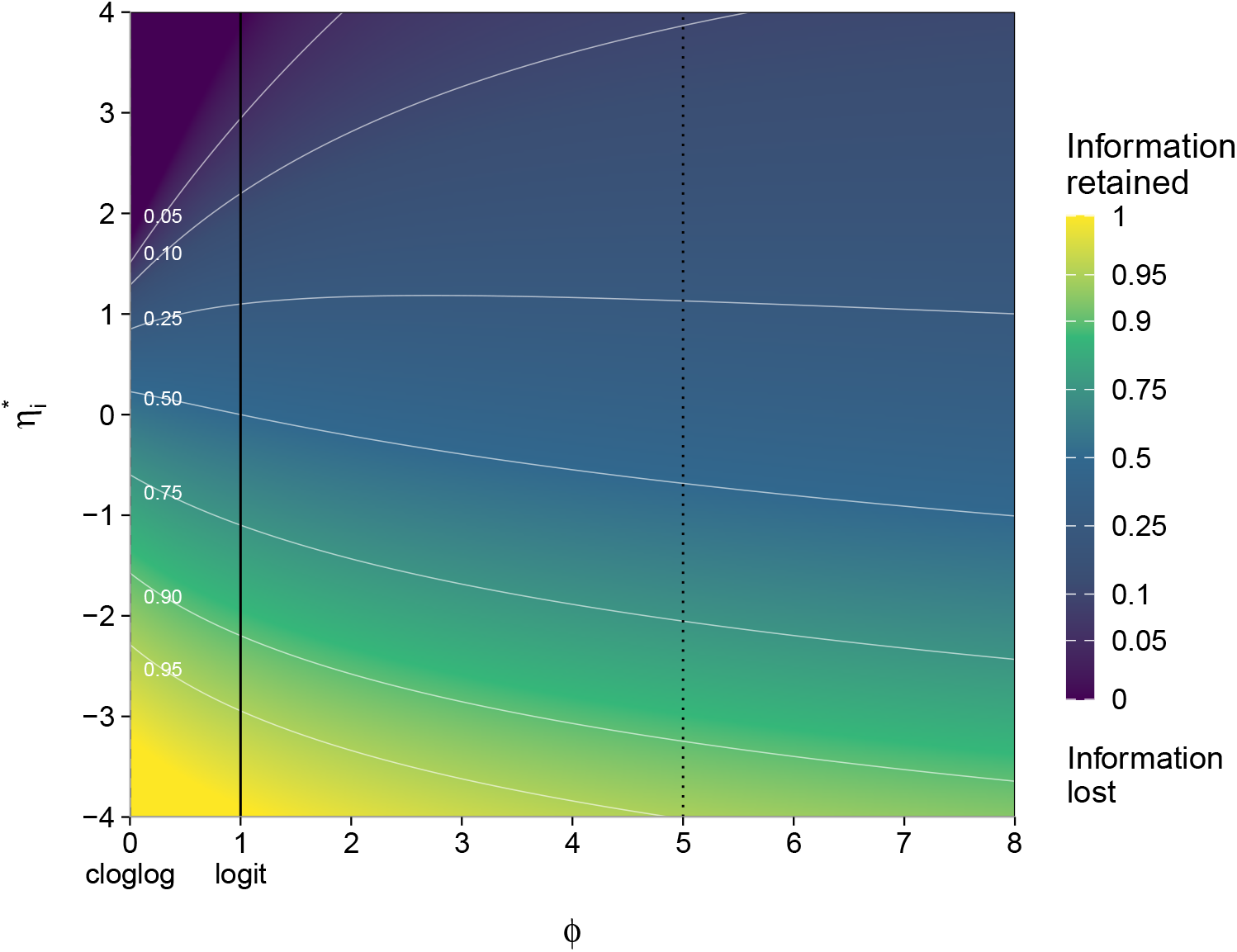
Information ratio as a bivariate function of the latent count linear predictor 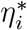 and the degree of overdispersion *ϕ*. At one, all information about the covariate effects forming 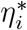 is retained in the gcloglog binomial regression, while at zero all information is lost. White contour lines indicate different proportions of information retained, ranging from 5% in the top (blue) part of the Figure to 95% in the bottom (yellow) part. The vertical slides corresponding to the case of equivariance and hence cloglog link, *ϕ* = 1 and hence logit link, and *ϕ* = 5 corresponding to more severe overdispersion.

### 3.2 Estimation of the shape parameter *ϕ* and sensitivity analysis

While more flexible, the use of the gcloglog link in practice requires estimation of the additional shape parameter *ϕ*. While maximum likelihood estimation jointly with the other parameters in the binomial regression (***β***) in equation (1) is possible, this is computationally burdensome and requires bespoke software implementations. As such, in the associated `gcloglog R`-package we opt for a simpler profiling approach based on the fact that, for fixed *ϕ*, the model reduces to a binomial GLM(M) albeit with a custom link function. This allows us to leverage existing, well-established software implementations that facilitate such model fitting, e.g., the `lme4 R`-package for estimating GLMMs (Bates et al., 2015, 2026).

Briefly, to profile the gcloglog link function we repeatedly fit the relevant binomial regression and maximize the outer likelihood with respect to *ϕ*. Such a profiling approach is similar to the method implemented for estimating the Tweedie power parameter *ξ* in the `tweedie R`-package (Dunn, 2026), as well as the estimator of the negative binomial shape parameter in the `MASS R`-package (Venables and Ripley, 2002). For GLMs, profiling is accomplished via a quasi-Newton routine for maximization as the first derivative of the likelihood with respect to *ϕ* is straightforward to obtain. For GLMMs among other more complex binomial regressions, we make use of derivative-free optimization, e.g., the `nloptr R`-package (Ypma and Johnson, 2025). After fitting, profile likelihood confidence intervals for the estimated 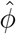 can be constructed based on adapting relevant functions in the `TMB R`-package (Kristensen et al., 2016).

In addition to estimating *ϕ*, the `gcloglog` package can facilitate sensitivity analysis to assess the severity of misspecifying the link function especially. For example, if we anticipate *ϕ* will be poorly estimated even under a larger sample size, then fitting gcloglog link binomial regression for increments of the shape parameter within the range of the profile confidence interval obtained above (especially if the intervals contains special cases such as *ϕ* = 1 or *ϕ ≈* 0) can be used to assess the stability of inference on the corresponding covariate effects. We demonstrate this in Section 5 on the two real data examples, where, due to estimation uncertainty in the shape of the link function, then the sign of parameter estimates for ***β*** can change within the confidence range of *ϕ*. In contrast, if the sign of parameter estimates and/or the exclusion of zero in the associated confidence intervals remains relatively stable across a range of *ϕ*, then the inferences on the covariate effects are likely to be robust across most plausible gcloglog link shapes.

## 4 Simulation study

We performed a numerical study to demonstrate the proposed gcloglog link, including estimation of the shape parameter *ϕ*, and the consequences of link misspecification when fitting a binomial regression. In the latter, we assess if fitting a cloglog or logit link, while the underlying link function tends more slowly to one (corresponding to *ϕ >* 1), exhibits similar consequences as ignoring overdispersion when fitting a Poisson regression model to true negative binomial distributed count data.

### 4.1 Use of the gcloglog link

We simulated Bernoulli and binomial (with *N*_*i*_ = 10 trials) responses from a binomial regression and proceeded to fit logit, cloglog, and gcloglog link binomial regressions to assess accuracy of the parameter estimates. For the last of the fitted models, we also tested whether the profile estimation approach proposed in Section 3.2 could recover the true link shape (or equivalently, the true *ϕ*).

In more detail, we examined two settings. The first setting simulated responses from a true model involving *n* = 500 observational units and covariates ***x***_*i*_ = (1, *x*_*i*_); *i* = 1, …, 500, where *x*_*i*_ was generated independently from a standard normal distribution. We set the corresponding vector of true coefficients to (*β*_0_ = 0, *β*_1_ = 2.3), and considered 100 possible values of *ϕ* in the range [0, 5]. For the second setting, we fixed *ϕ* = 0 or 1, i.e., the true link function was either the cloglog or logit link, and simulated 1000 datasets at varying sample size 50 *≤ n ≤* 600. The rest of this setup was the same as the first setting.

Figure 3 presents results from the first setting, i.e., with fixed *n* and varying *ϕ*. Since estimation of the additional shape parameter in the gcloglog link exhibited considerable variability especially at large *ϕ*, we chose to exclude extreme simulation results for covariate estimates *β* and link shape *ϕ*, namely those that were more than 40 median absolute deviations from the median. Across all settings of true *ϕ*, gcloglog link binomial regression performed reasonably well at recovering the covariate effects, while the cloglog and logit links unsurprisingly exhibited noticeable bias when the true *ϕ* deviated far away from *ϕ* = 0 and *ϕ* = 1 respectively. On the other hand, the reasonable performance of the gcloglog link came at the cost of substantially increased estimation variability, as reflected by the 2.5% and 95.5% percentiles for parameter estimates, although this variability decreased substantially when moving from Bernoulli to binomial *N*_*i*_ = 10 responses. From the third row in Figure 3, we see that when the true *ϕ* was reasonably small, including the cases of 0 and 1 corresponding to cloglog and logit links respectively, the profile estimation approach of Section 3.2 for the gcloglog link estimated *ϕ* well. Performance deteriorated though when *ϕ* became larger, with consistent overestimation in the case of Bernoulli responses, and to a much lesser extent with binomial *N*_*i*_ = 10 responses.

**Figure 3:**
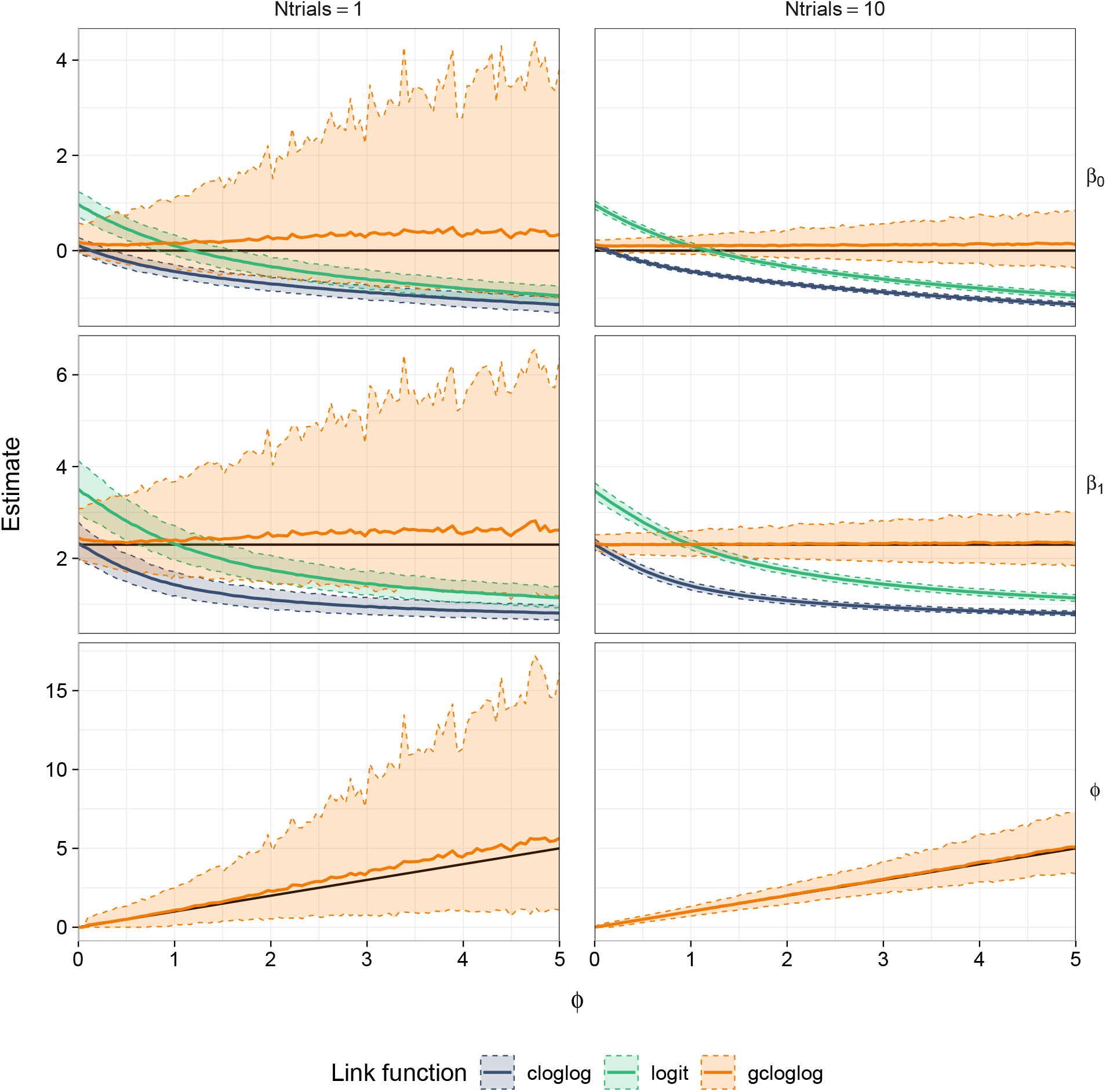
Results from the first setting of simulations in Section 4.1, involving fixed *n* = 500 and varying the shape parameter 0 ≤ *ϕ* ≤ 5. In panel (a), the two columns correspond to Bernoulli responses and binomial *N*_*i*_ = 10 responses, respectively. The mean estimated co-efficients (averaged across 1000 datasets for each value of *ϕ*) are shown as solid lines, along with colored ribbons corresponding to 2.5% and 97.5% percentile range parameter estimates, from fitted binomial regressions are shown, assuming either the cloglog (blue), logit (green), or gcloglog (orange) link functions. The value of the true parameter is indicated by a black solid line. In panel (b), the mean estimated *ϕ* is shown for Bernoulli and binomial responses.

Figure 4 presents results from the second setting, i.e., with fixed *ϕ* and varying *n*. As expected, the variability of the parameter estimates from gcloglog link binomial regression fits unsurprisingly decreased with increasing *n*, but was substantially larger compared with the variability that comes from fixing *ϕ* at its true value. Indeed, for Bernoulli responses the proportion of gcloglog link regression fits requiring exclusion of outliers (more than 40 median absolute deviations from the median, as described above) was negligible for *ϕ* near 0 or 1, but reached approximately 15% at *ϕ* = 5 and 24% at *n* = 50, while reaching below one percent at *n ≈* 300. For binomial responses with *N*_*i*_ = 10 trials, no simulated dataset required exclusion for any value of *ϕ* and *n* when fitting the gcloglog link, although the variability is still noticeably larger compared with the fixed *ϕ* link regressions.

**Figure 4:**
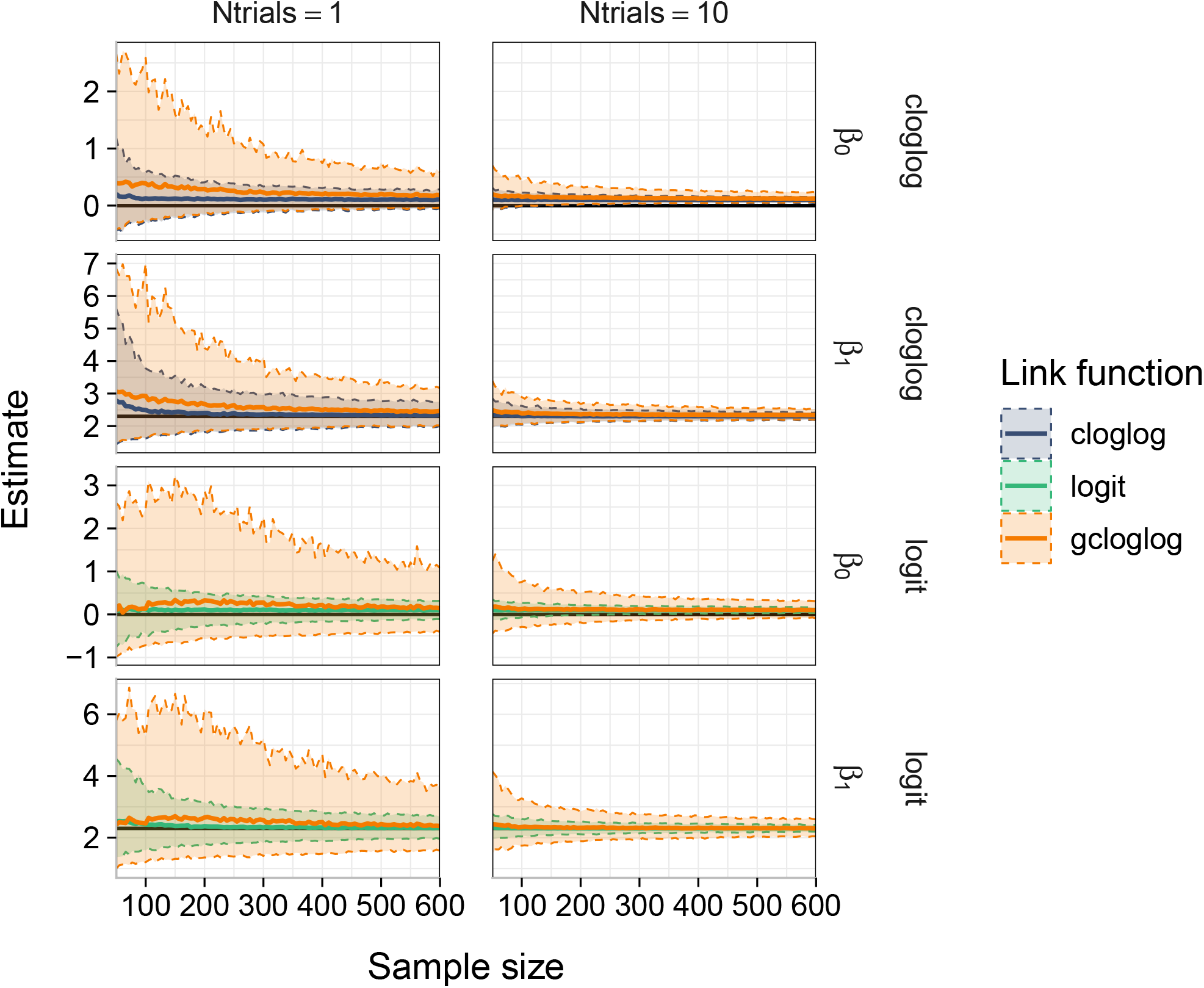
Results from the second setting of simulations in Section 4.1, involving fixed *ϕ* = 0 (cloglog link; first and third rows) and 1 (logit link; second and fourth rows) and varying sample size *n*. The two columns correspond to Bernoulli responses and binomial *N*_*i*_ = 10 responses, respectively. The mean estimated coefficients (averaged across 1000 datasets for each value of *n*) are shown as solid lines, along with colored ribbons corresponding to 2.5% and 97.5% percentile range parameter estimates, from fitted binomial regressions are shown, assuming either the cloglog (blue), logit (green), or gcloglog (orange) link functions. The value of the true parameter is indicated by a black solid line.

### 4.2 Uncertainty estimates

In this simulation, we explored the consequences of link misspecification in binomial regression. To this end, we simulated 1000 datasets each of which comprised *n* = 500 Bernoulli responses generated from a gcloglog link binomial regression, with covariates (1, *x*_*i*1_, *x*_*i*2_; *i* = 1, …, *n* where *x*_*i*1_ and *x*_*i*2_ were independently simulated from a standard normal distribution, and where the true coefficient values were set to (*β*_0_ = 0, *β*_1_ = 2.3, *β*_2_ = 0). Similar to the first setting in Section 4.1, the true shape parameter was varied across 100 true values of 0 *≤ ϕ ≤* 5. For each simulated dataset, we fitted binomial regression assuming a cloglog, logit, or a gcloglog link function.

Next, to construct standard errors for the coefficient estimates, a standard approach is to use the inverse of the expected information matrix from the fitted GLM, e.g., via the `vcov` function as applied to the fitted `glm` object in `R`. While this is fine for the cloglog and logit links, for gclogclog link such standard errors ignore the variability due to the estimation of the shape parameter *ϕ*, and consequently are not expected to produce nominal coverage. To correct for this, we applied a correction based on the joint information; see Appendix S2 for further details of this. Another, arguably more naive but practical approach is to use sandwich-based standard errors, e.g., via the `sandwich` package Zeileis 2006, which provide yet another attempt to account for model misspecification when computing standard errors. On the other hand, since link function misspecification does not directly equate to misspecification of the variance function, then we do not necessarily expect these to work well either.

The results of the simulations are displayed in Figure 5a (standard errors) and Figure 5b (coverage). As expected, standard errors of gclogclog link binomial regression based on the joint information provided good coverage, while naive standard errors that ignored extra estimation variability due to the shape parameter provided poor coverage. Sandwich-based standard errors did not provide improved coverage for cloglog and logit regression. The cloglog and logit link binomial regressions provided a poor estimate for the standard error across *ϕ*, with the obvious exception of *ϕ* = 0 and *ϕ* = 1, respectively, for both ordinary and sandwhich-based standard errors. As a result of the underestimation, as well as due to the downward bias displayed in Figure 3, coverage of the confidence intervals for both models decreased with increasing overdispersion in the latent count, and distance between their asssumed *ϕ* and the true *ϕ*. Regardless of the link function, coverage for the absence of an effect, as in the case of *β*_2_, was well estimated.

**Figure 5:**
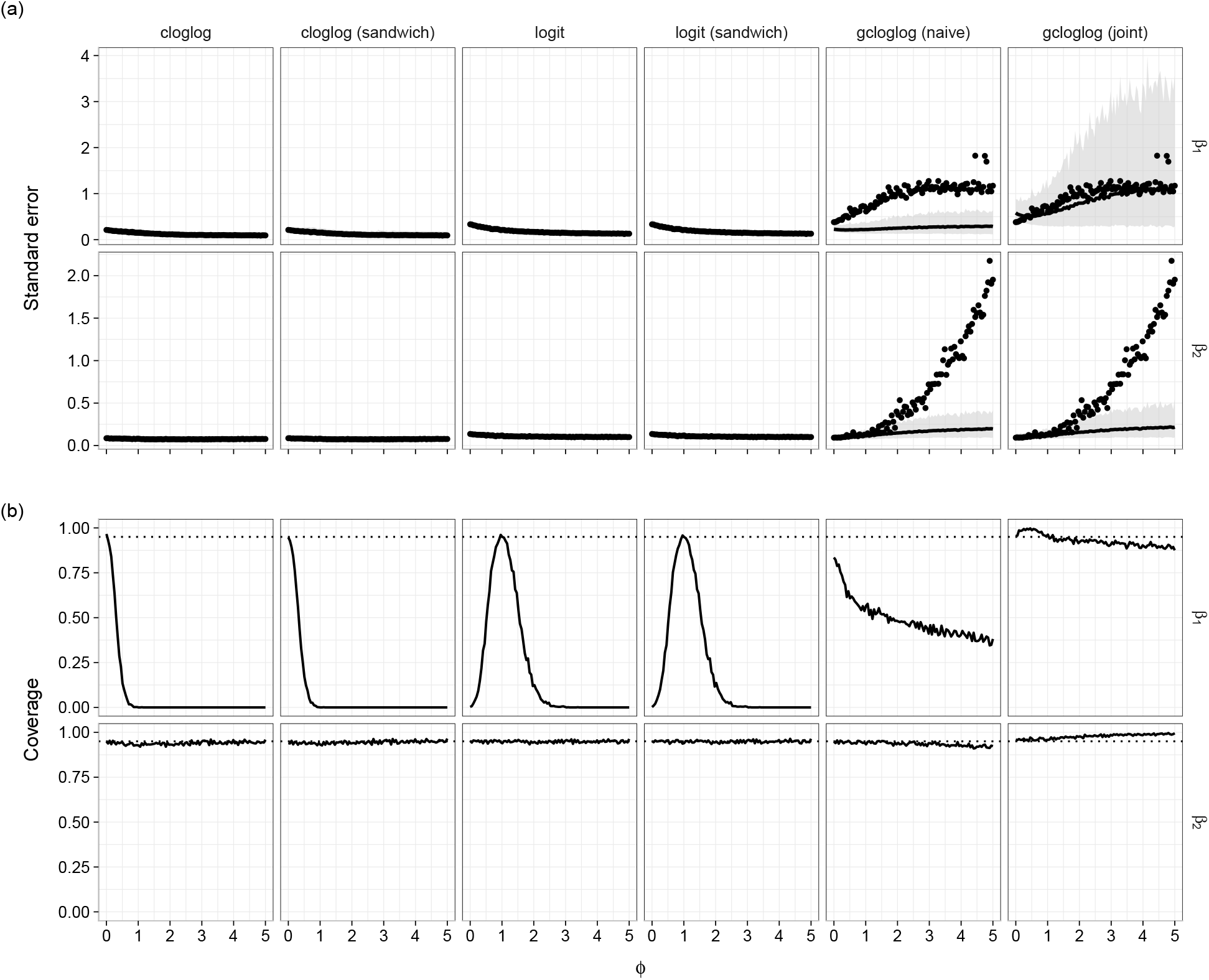
Results from the simulation in Section 4.2, involving fixed *n* = 500 and varying the shape parameter 0 ≤ *ϕ* ≤ 5. The true model simulated Bernoulli responses from a gcloglog link binomial regression for a given *ϕ*, with two slope parameters with *β*_1_ = 2.3 (top row of each panel) and *β*_2_ = 0 (bottom row of each panel). For each simulated dataset, we fitted binomial regression assuming either the cloglog (leftmost two columns), logit (middle two columns) link, and the gcloglog (rightmost two columns) link functions. For the cloglog and logit models, the ordinary standard error and sandwich-based standard error are shown. For the gcloglog model, the naive joint standard errors are shown. In panel (a), the mean standard errors (averaged across 1000 simulated datasets) is shown as a solid black line, with the grey ribbon corresponding to 2.5% and 97.5% percentile ranges In panel (b), coverage of the true parameter by the confidence interval is displayed by a dashed black line, and nominal 95% coverage is displayed as a dotted line.

## 5 Case studies

We demonstrate the use of gcloglog link binomial regression, along with the idea of performing sensitivity analysis around the choice of link function, on two real ecological datasets: 1) Bat acoustic monitoring data (Cole, 2022), and 2) Salamander count data (Price et al., 2016, 2015). In both cases, binomial regressions were fitted using the `lme4 R`-package (Bates et al., 2026) for its ability to fit binomial models with custom link functions. On the other hand, negative binomial models were fitted with the `glmmTMB R`-package (Brooks et al., 2017), as `lme4` does not return confidence intervals for the negative binomial shape parameter.

### 5.1 Bat data

The Bat monitoring data was collected by Cole (2022) and is available as part of the `EcoCountHelper R`-package (Cole et al., 2022). Briefly, the data were collected in the summers of 2016 and 2017 in Grand Teton National Park (US) by conducting acoustic monitoring of bat call sequences at 40 sites and for seven different species. Here, we restrict our analysis to a single species, namely the big brown bat (*Eptesicus fuscus*). The data also includes various covariates including: (i) the inverse distance to porous buildings surrounding the site, (ii) the proportion of lights that had a cool hue, (iii) brightness score of lights in the vicinity, (iv) the year of data collection, (v) the ordinal day of data collection, (vi) elevation, (vii) the distance to the nearest water source, (viii) the proportion of developed land surrounding the site, (ix) the proportion of forest surrounding the site, and (x) the moon’s fraction that was illuminated

We fitted four GLMMs, all of which contained the aforementioned covariates as fixed effects along with site as a random intercept. Specifically, as a benchmark we fitted a log-link NBII GLMM to the original count responses, and then truncated the counts to Bernoulli responses before fitting binomial GLMMs assuming either the cloglog, logit, and gcloglog link functions.

Figure 6a presents a comparison of the fixed effect estimates 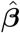 along with corresponding 95% confidence intervals. Overall, all four models produced fairly similar trends, for example, offering statistically clear evidence of a positive effect for (inverse) distance to porous buildings and the day the data were collected. The cloglog link binomial GLMM produced the most different estimates from the benchmark negative binomial GLMM, while its associated confidence intervals were narrower compared to its logit and gcloglog link counterparts. Indeed, gcloglog link binomial GLMM provided the closest match to the negative binomial GLMM, with the root mean squared error (RMSE) between the two sets of parameter estimates equaling 0.210, compared with 0.236 for the logit link and 0.288 for the cloglog link. On the other hand, for some covariates such as year of data collection, the negative binomial regression produced a smaller confidence interval leading to statistical significance, while the corresponding intervals for the three binomial regressions were wider and contained zero. This is consistent with the discussion in Section 3.1 on information loss, and indeed for all fixed effects the confidence intervals were consistently shorter for the negative binomial GLMM for count data compared with the binomial GLMMs for Bernoulli data.

**Figure 6:**
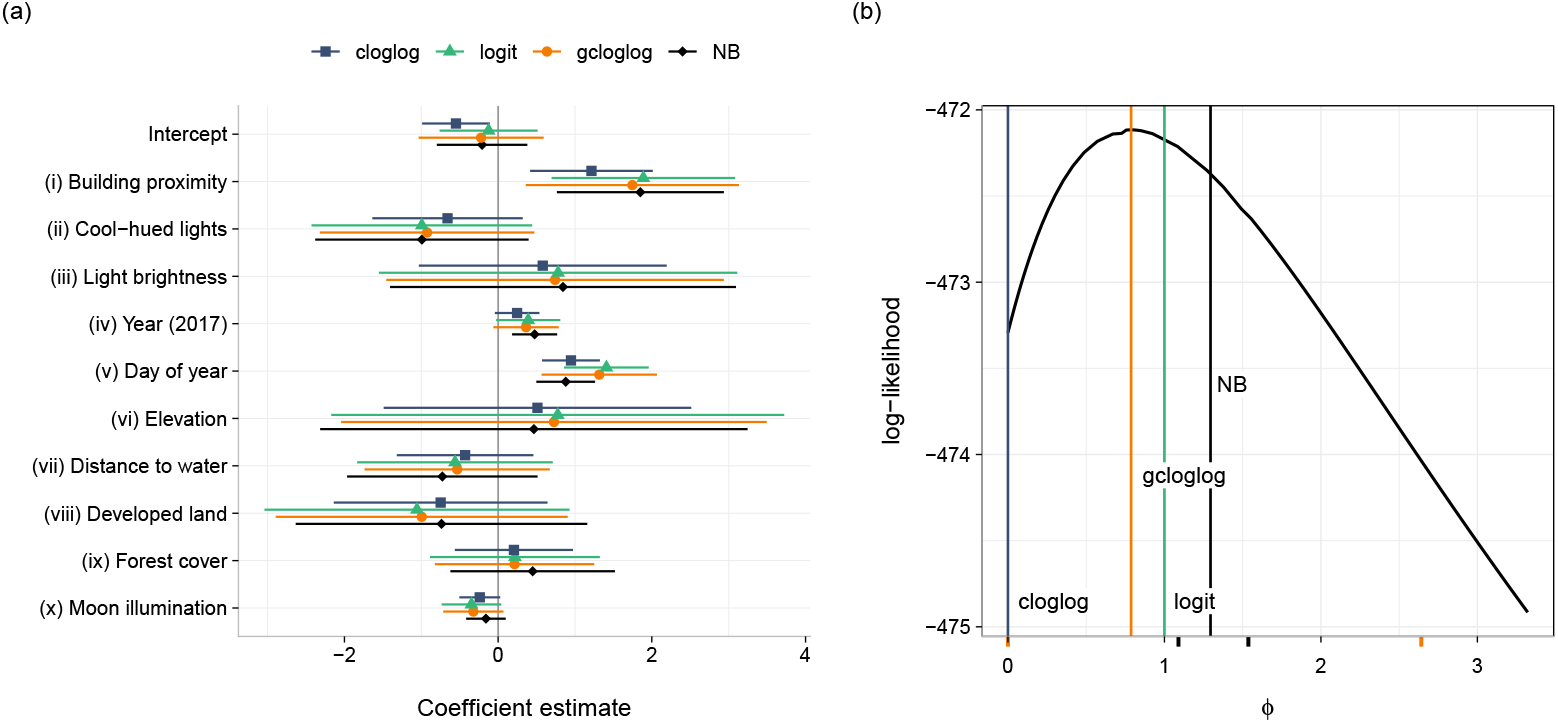
Panel (a) Fixed-effect estimates and corresponding 95% confidence intervals for application to the Bat data, comparing a negative binomial (NB) GLMM fitted to the original count data, with binomial GLMMs fitted to the truncated Bernoulli responses assuming a gcloglog, logit, and cloglog link functions. Panel (b) presents the profile likelihood for the shape parameter *ϕ* in the gcloglog link GLMM. The orange vertical line marks the estimate of 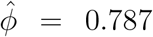 (with a corresponding 95% CI of (0,2.642) shown as orange tick marks). Vertical lines mark the logit (green; *ϕ* = 1) and cloglog (blue; *ϕ* = 0) link binomial models, as well as the negative binomial GLMM estimate (black) *ϕ* = 1.295 (with a corresponding 95% CI (1.091, 1.538) shown as green tick marks).

Turning to the other model parameters, the estimated variance for the site-level random intercept was 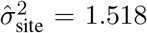 for the negative binomial GLMM, versus 1.525 for the logit, 1.315 for the gcloglog, and 0.694 for the cloglog binomial models. That is, the logit link binomial GLMM was closest to the negative binomial estimate when it came to the variance component, while the cloglog link model estimated the smallest site-level heterogeneity of the four GLMMs compared. With regards to the dispersion parameter, Figure 6b presents the profile likelihood for *ϕ* for the fitted gcloglog model, with the logit (*ϕ* = 1), cloglog (*ϕ* = 0), negative binomial (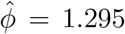 with a corresponding 95% CI of 1.091 to 1.538) GLMMs also included for reference. Profile estimation for the gcloglog link binomial GLMM produced a maximum likelihood estimate of 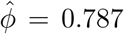 along with a corresponding 95% confidence interval ranging (0, 2.642). Note the latter includes the estimate and confidence interval from the negative binomial GLMM, as well as binomial GLMMs assuming the cloglog (*ϕ →* 0) and logit (*ϕ* = 1) link functions, but is not contained in the confidence interval of the shape parameter from the negative binomial.

To conclude our application, we performed a sensitivity analysis based on fitting gcloglog link binomial GLMMs for a grid of values of *ϕ* that span the range of its estimated 95% confidence interval given above. The results of this are presented in Appendix S3, which, concordant with Figure 6a, show that the confidence intervals for all ten slope coefficients as well as the intercept were consistent in terms of including or excluding zero. This suggested no potential change and hence a reasonable degree of robustness in the inference of the gcloglog link model across a plausible set of values for true *ϕ*. Additionally, we examined how the aforementioned RMSE to the fixed effects estimates from the NB GLMM varied as a function of *ϕ*, although we acknowledge that in practice this is not possible if the count data were not available. In summary, the RMSE produced by the gcloglog model was lower than that of the other two binomial models, and although it did not correspond to the lowest RMSE for all evaluated *ϕ* which was located at approximately *ϕ* = 0.5. The RMSE declined from *ϕ* = 0 to the minimum, rising slowly to the profiled solution at *ϕ* = 0.787, and increasing more sharply for larger values of *ϕ*. Interestingly, near the negative binomial GLMM estimate of *ϕ* = 1.295, the RMSE had increased to about 0.25, and was thus further removed from the minimum RMSE of 0.2 reached by the estimate of the gcloglog model.

### 5.2 Salamander data

The Salamander data were collected by Price et al. (2016), and are available via the `glmmTMB R`-package (Brooks et al., 2017). Briefly, counts of stream salamanders were recorded at 23 streams in south-eastern Kentucky, United States. Each creek was sampled four times. Along with the count responses, covariates recorded include: (i-vi) species-life stage, which was a factor variable including *G. porphyriticus* as the reference level, (i) *P. ruber*, (ii) *D. monticola*, adults (iii) and larvae (iv) of *E. cirrigera*, (v) larvae of *Desmognathus* and (vi) *D. fuscus*, (vii) a binary factor indicating whether a site had been affected by mountaintop-removal coal mining, with “mined” as the reference level, (viii) the amount of cover objects in the stream, (ix) days since the last precipitation, and (x) day of year. Analogous to the previous example, we began by fitting a NBII GLMM to the count response as a benchmark, including all covariates above as fixed effects along with a random intercept to account for replication at each streams. The resulting fit estimated a dispersion parameter of 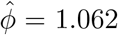 (with a corresponding 95% CI of 0.809 to 1.376), which suggested that a logit link binomial regression model may be reasonable when analyzing the associated truncated Bernoulli responses.

Figure 7a presents the fixed effect estimates and corresponding 95% confidence intervals for the four models of interest, i.e., the NBII GLMM along with the cloglog, logit, and gcloglog link GLMMs. Again, the overall conclusions for the drivers of Salamander data were fairly similar across the four GLMM fits e.g., there was statistical evidence of lower mean (latent) abundance for some species and life stages (e.g., *P. ruber* and *E. cirrigera* adults) compared to the reference (*G. porphyriticus*), along with mountaintop-removal coal mining (noting the reference category is “mined”). Interestingly, the fixed effect estimates from the gcloglog link GLMM were the furthest from the negative binomial GLMM, with an RMSE between the two sets of parameter vectors equaling 0.450, compared with the logit link (RMSE = 0.294) and cloglog link (RMSE = 0.226) models. Similarly, the estimated variance component for stream from the gcloglog link GLMM was nearly triple that of the negative binomial GLMM estimate (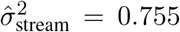 versus 0.277); the corresponding estimates for the logit and cloglog link binomial GLMMs were 0.569 and 0.297, respectively. These results are despite the estimate value of *ϕ* from the negative binomial GLMM corresponding closest to the logit (and not the cloglog) link. Finally, fitting a gcloglog link binomial GLMM produced a larger estimate of the shape parameter 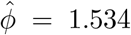 compared with the negative binomial GLMM and the confidence interval it produced for the shape parameter (Figure 7b). On the other hand, its corresponding 95% confidence interval was fairly wide at (0, 3.79), thus containing the negative binomial shape parameter estimate and its confidence interval. As a result this covered all three other models’ values of *ϕ*.

**Figure 7:**
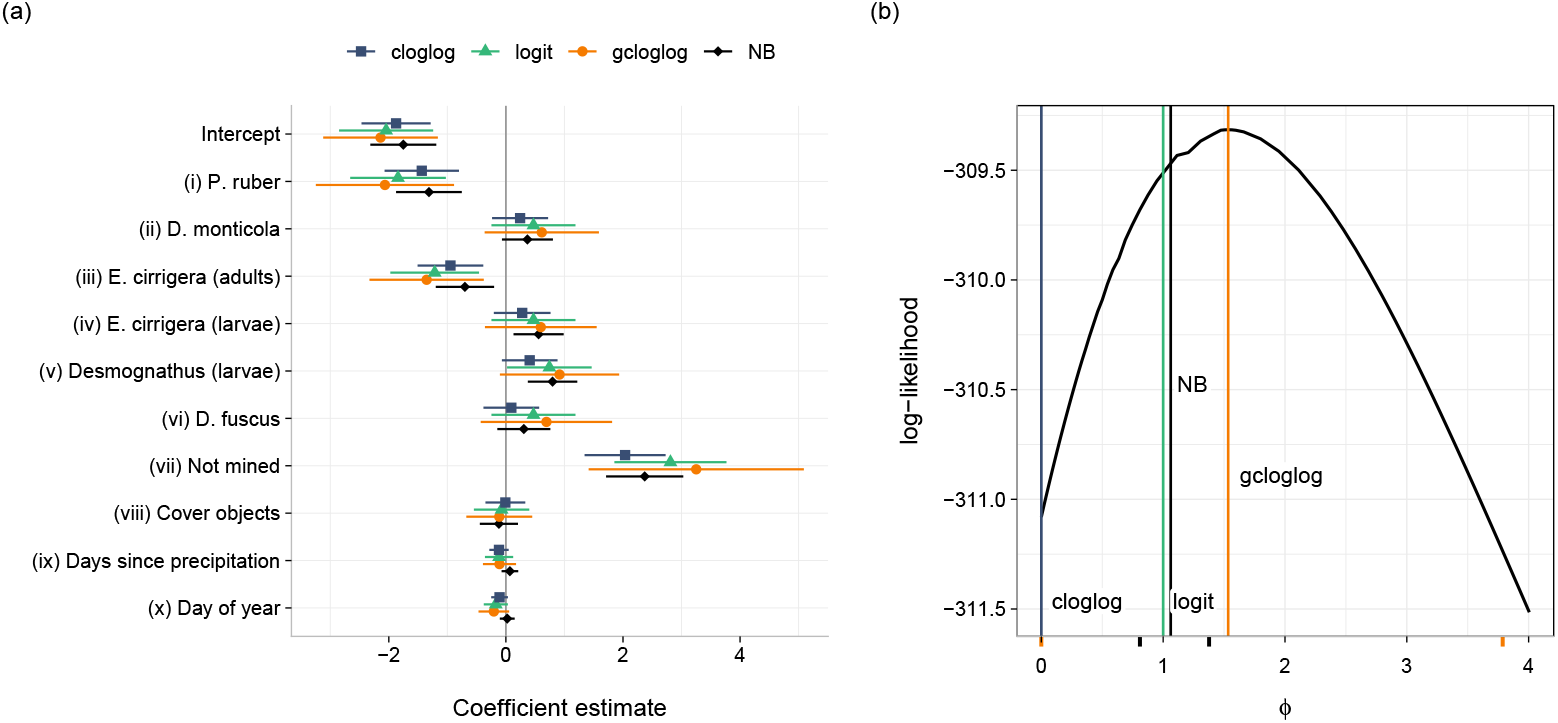
(a) Fixed-effect estimates and corresponding 95% confidence intervals for application to the Salamander data, comparing a negative binomial (NB) GLMM fitted to the original count data, with binomial GLMMs fitted to the truncated Bernoulli responses assuming a gcloglog, logit, and cloglog link functions. Panel (b) presents the profile likelihood for the shape parameter *ϕ* in the gcloglog link GLMM. The orange line marks the estimate of 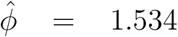 (with a corresponding 95% CI (0, 3.787) shown as orange tick marks). Vertical lines mark the the logit (green; *ϕ* = 1) and cloglog (blue; *ϕ* = 0) link binomial models, as well as the negative binomial GLMM estimate (black) of *ϕ* = 1.064 (with a corresponding 95% CI (0.809, 1.376) shown as green tick marks).

In Appendix S3 we present results from a sensitivity analysis of the gcloglog link binomial model (S3). For most fixed effects, inference did not change with the same sign and statistical significance was retained across a plausible range of values of true *ϕ*. We also examined how the RMSE to the estimates from the NB GLMM (see appendix S4)varied as a function of *ϕ*, although we acknowledge that in practice this is not possible if the count data were not available. Interestingly, none of the three binomial GLMMs produced the minimum RMSE to the fixed effect estimates from the negative binomial model. Moreover, the RMSE was asymmetric as it declined only slightly from *ϕ* = 0 to a shallow minimum around *ϕ ≈* 0.4, before increased steadily reaching the RMSE of 0.450 at the profiled maximum likelihood estimate from the gcloglog link binomial GLMM of 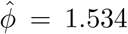, and an RMSE of 0.601 by *ϕ* = 2. Consistent with the above results, the value of *ϕ* that maximized the profile likelihood was far removed from the value that minimized RMSE to the negative binomial model: this is again not unexpected given the limited number of replications.

## 6 Discussion

In ecology, binary and binomial data, such as the presence or absence of a species at a site, often arise from the truncation (whether through the sampling process or subsequent data wrangling) of an underlying measure of abundance. As such, when developing a binomial regression for such data it is important to keep in mind that the choice of link function represents assumptions about the latent abundance process.

In this article, we showed that if the latent abundance process follows a distribution in the Tweedie class, and the mean is modeled, then binomial regression with a cloglog link function does not generally provide directly interpretable parameters. The exception to this is the Poisson distribution (see also the Poisson-Gamma formulation of the Tweedie distribution, Foster and Bravington, 2013). If the latent abundance process follows a negative binomial distribution with linear mean-variance function, then cloglog link binomial regression can recover the slope effects only. To handle the more common and arguably more relevant case where the latent abundance process follows a negative binomial distribution with quadratic mean-variance relationship, we derived the generalized complementary log-log link function, which involves an additional shape parameter and includes the cloglog and logit links as special cases. The case of the logit link is especially interesting, and we show that it can arise from the truncation of a latent abundance process that is negative binomial distributed with moderate overdispersion (*ϕ* = 1).

Through simulations and two case studies, we demonstrated that estimation of the shape parameter in the gcloglog link is challenging and requires replication for accurate estimation. However, the simulations also emphasized its potential efficacy with sufficiently large sample sizes, particularly when compared to the case of link misspecification, i.e., selecting a link function in binomial regression that misrepresents the latent abundance process. The latter can result in biased parameter estimates and invalid inference, e.g., if a cloglog link is used when the latent count process is overdispersed, analogous to the anticonservative that results when fitting a Poisson distribution to overdispersed counts (Harrison, 2014). This was also seen to some extent with the Bat data case study, where cloglog link binomial regression produced the most different fixed effect estimates and inference compared with its logit and gcloglog link counterparts. Conversely, for the Salamander data case study the smaller sample size led to gcloglog link binomial producing the most different estimates to the benchmark negative binomial regression.

It is important to emphasize that the accompanying `gcloglog R`-package serves a dual purposes of fitting a gcloglog link binomial regression (including profile estimation of *ϕ*) to provide inference on the underlying latent count process, and acting as a diagnostic tool for sensitivity analysis to assess the stability of inference when the link (synonymous to the degree of overdispersion in the latent process) is misspecified. Indeed, we stress that the latter is more potentially important: the gcloglog link should not be viewed as a panacea let alone a default link in binomial regression for ecologists. Rather, it provides a basis for practitioners to more explicitly think about the link specification of binomial regression in ecological datasets, balancing interpretability and computational convenience with assumed properties of the latent abundance process.

For binomial data, the additional flexibility accommodated by the gcloglog link function is distinct from the overdispersion modeled by specifying a beta-binomial distribution (Crowder, 1978). A gcloglog model modifies the link function to accommodate overdispersion in the latent abundance process by bending the link function, while beta-binomial models accommodate unobserved heterogeneity in the probability of success between clusters by augmenting the variance function (Harrison, 2015; Crowder, 1978; Skellam, 1948). Consequently, the beta-binomial accommodates a different kind of overdispersion, namely acting on the response directly and relative to the binomial variance function. However, the mean specification and chosen link function remain the same as in a binomial regression. More generally, while this article has focused on explicit connections with biomass and count responses, the developments can straightforwardly be extended to other types of abundance data such as percent cover in the form of ordinal classes or continuous cover measurements (Kubinec, 2023; Korhonen et al., 2024), and zero-inflated response distributions (Wenger and Freeman, 2008; Zuur et al., 2009). The latter, especially if the probability of excess zeros is also modeled, are likely to lead to even more complex link functions and results than those seen in this article, as the probability of a positive response then depends on both the mean models of the excess zeros and of the non-zero distribution.

## Acknowledgments

Francis Hui was supported by an Australian Research Council Discovery Project DP240100143.

## Data availability statement

No new data were collected for this study. The bat acoustic monitoring data are available in the `EcoCountHelper R`-package (Cole et al., 2022), and the salamander data are available in the `glmmTMB R`-package (Brooks et al., 2017; Price et al., 2015).

