## Supplementary information for "On Motivating the Choice of Link Function in Binomial Regression Through A Latent Abundance Process"

### S1 Derivation of information ratios

#### S1.1 Underlying Poisson count

Consider a set of $N$ independent observations $(x_i, y_i^*); i = 1, \dots, N$ where $y^*$ follows a log-link Poisson regression. That is, $y_i^* \sim$ Poisson$(\mu_i)$ with $\log(\mu_i) = \beta_0 + x_i\beta_1$. It is straightforward then to show that the gradient is

$$\nabla_{\mu_i}\ell_{\text{Pois}} = \frac{y_i^*}{\mu_i} - 1,$$

and thus its expected information given by

$$\mathcal{I}_{\text{Pois}}(\mu_i) = -\mathbb{E}\left(\frac{\partial \nabla_{\mu_i}\ell_{\text{Pois}}}{\partial \mu_i}\right) = \frac{\mathbb{E}(y_i^*)}{\mu_i^2} = \mu_i^{-1}.$$

After truncating the counts such that $y_i = I(y_i^* > 0)$ and fitting a cloglog link binary regression, i.e., $y_i \sim$ Bernoulli$(p_i)$ with $\log(-\log(1 - p_i)) = \eta_i$, by definition we have the log-likelihood

$$\ell_{\text{Bern}}^{\text{cloglog}} = \sum_{i=1}^{N} y_i \log(p_i) + (1 - y_i)\log(1 - p_i) = \sum_{i=1}^{N} y_i \log(1 - \exp(-\mu_i)) - (1 - y_i)\mu_i.$$

Consequently, it follows that

$$\nabla_{\mu_i}\ell_{\text{Bern}}^{\text{cloglog}} = y_i \frac{\exp(-\mu_i)}{1 - \exp(-\mu_i)} - (1 - y_i) = y_i \frac{1}{\exp(\mu_i) - 1} - (1 - y_i)$$

and thus

$$\mathcal{I}_{\text{Bern}}^{\text{cloglog}}(\mu_i) = -\mathbb{E}\left(\frac{\partial \nabla(\mu_i)\ell_{\text{Bern}}^{\text{cloglog}}}{\partial \mu_i}\right) = \frac{1 - \exp(-\mu_i)}{\exp(-\mu_i) + \exp(\mu_i) - 2} = \frac{1}{\exp(\mu_i) - 1}.$$

It follows that the per-observation "information ratio" is given by

$$\text{IR}_i = \frac{\mathcal{I}_{\text{Bern}}^{\text{cloglog}}(\mu_i)}{\mathcal{I}_{\text{Pois}}(\mu_i)} = \frac{\mu_i}{(\exp(\mu_i) - 1)}.$$

This ratio is strictly between 0 and 1, monotonically decreasing and tends to one as $\mu_i \to 0$, and tends to zero as $\mu_i \to \infty$.

#### S1.2 Negative binomial count with $\phi = 1$

Consider a set of $N$ independent observations where $y^*$ follows a log link, quadratic mean-variance negative binomial regression with $\phi = 1$ i.e., Geometric regression. That is, $y_i^* \sim \text{Geometric}(\mu_i)$ and $\log(\mu_i) = \eta_i$. The associated log-likelihood function is then given by

$$\ell_{\text{Geom}} = \sum_{i=1}^{N} y_i^* \log(\mu_i) - (y_i^* + 1)\log(1 + \mu_i),$$

so that, following the same algebraic derivation as for the Poisson count case, the expected information is given by

$$\mathcal{I}_{\text{Geom}}(\mu_i) = \mu_i^{-1} + \frac{1}{1 + \mu_i} = \frac{1}{\mu_i(1 + \mu_i)}.$$

Suppose now we truncate the counts such that $y_i = I(y_i^* > 0)$ and fit the logit link binary regression, i.e., $y_i \sim \text{Bernoulli}(p_i)$ with $\log\left(\frac{p_i}{1-p_i}\right) = \eta_i$. Straightforward algebra then reveals that the expected information is given by

$$\mathcal{I}_{\text{Bern}}^{\text{logit}}(\mu_i) = \frac{1}{\mu_i(1 + \mu_i)^2}.$$

The per-observation "information ratio" is thus given by

$$\text{IR}_i = \frac{\mathcal{I}_{\text{Bern}}^{logit}(\mu_i)}{\mathcal{I}_{\text{Geom}}(\mu_i)} = \frac{1}{1 + \mu_i}.$$

This ratio is again strictly between 0 and 1, monotonically decreasing and tends to one as $\mu_i \to 0$, and tends to zero as $\mu_i \to \infty$.

Note as $\mu \to \infty$, the information ratio for the geometric case tends to zero much slower than for the Poisson case, i.e., polynomial versus exponential decay. This is because the probability of positive count tends to one much quicker for a Poisson distribution $(1 - \exp(-\mu_i))$ compared to the geometric distribution $1/(1 + \mu_i)$, meaning the information loss due to truncation is much quicker for the former.

#### S1.3 Negative binomial count with $\phi \in (0, \infty)$

Consider a set of $N$ independent observations where $y^*$ follows a log link, quadratic mean-variance negative binomial regression with general $\phi$. That is, $y_i^* \sim \text{NBII}(\mu_i, \phi)$ with $\log(\mu_i) = \eta_i$. Then the corresponding log-likelihood function, ignoring constants, is given by

$$\ell_{\text{NB}} = \sum_{i=1}^{N} \log\Gamma(y_i^* + \phi^{-1}) - \log\Gamma(\phi^{-1}) + \phi^{-1}\log(\phi^{-1}) + y_i^* \log(\mu_i) - (y_i^* + \phi^{-1})\log(\mu_i + \phi^{-1}).$$

Following similar steps similar to the above, the expected information is then given by

$$\mathcal{I}_{\text{NB}}(\mu_i) = \frac{\phi^{-1}}{\mu_i(\mu_i + \phi^{-1})} = \frac{1}{\mu(\mu\phi + 1)}.$$

Suppose now we truncate the counts such that $y_i = I(y_i^* > 0)$ and fit a gcloglog link binary regression, i.e.,

$$y_i \sim \text{Bernoulli}(p_i), \quad \log\left(\frac{\phi^{-1}}{(1-p_i)^{\phi}} - \frac{1}{\phi}\right) = \eta_i.$$

The leads to the corresponding log-likelihood function

$$\ell_{\text{Bern}}^{\text{gcloglog}} = \sum_{i=1}^{N} y_i \log(p_i) + (1-y_i)\log(1-p_i) = \sum_{i=1}^{N} y_i \log(1 - (1+\phi\mu_i)^{-\phi^{-1}}) - \frac{(1-y_i)}{\phi}\log(1+\phi\mu_i).$$

Therefore, we obtain the corresponding expected information

$$\mathcal{I}_{\text{Bern}}^{\text{gcloglog}}(\mu_i) = \frac{1}{(1+\phi\mu_i)^2((1+\phi\mu_i)^{\frac{1}{\phi}} - 1)},$$

and the resulting per-observation "information ratio" is given by

$$\text{IR}_i = \frac{\mathcal{I}_{\text{Bern}}^{\text{gcloglog}}(\mu_i)}{\mathcal{I}_{\text{NB}}(\mu_i)} = \frac{\mu_i}{(1+\phi\mu_i)((1+\phi\mu_i)^{\phi^{-1}} - 1)}.$$

This ratio thus covers the general case, including both the Poisson/cloglog and geometric/logit as special cases for $\phi = 0$ and $\phi = 1$, respectively.

### S2 Joint information for gcloglog link binomial regression

Note that

$$I_{\text{Bern}}^{\text{gcloglog}} = \begin{bmatrix} \mathcal{I}_{\log\phi,\log\phi} & \mathcal{I}_{\log\phi,\beta} \\ \mathcal{I}_{\beta,\log\phi} & \mathcal{I}_{\beta,\beta} \end{bmatrix},$$

where $I_{\beta,\beta}$ denotes the observed information of a GLM fitted conditional on $\phi$. In order to calculate confidence intervals for a gcloglog fit binomial regression then, the other two matrices need to be separately derived.

We have the gcloglog binomial log-likelihood function

$$\ell_{\text{Bern}}^{\text{gcloglog}} = \sum_{i=1}^{N} y_i \log(p_i) + (1-y_i)\log(1-p_i) = \sum_{i=1}^{N} y_i \log(1-(1+\phi\mu_i)^{-\phi^{-1}}) - \frac{(1-y_i)}{\phi}\log(1+\phi\mu_i),$$

so that

$$\frac{\partial \ell_{\text{Bern}}^{\text{gcloglog}}}{\partial \log\phi} = \sum_{i=1}^{n} \frac{\partial \ell_{\text{Bern}}^{\text{gcloglog}}}{\partial p_i}\frac{\partial p_i}{\partial \phi}\frac{\partial \phi}{\partial \log\phi}.$$

The first partial derivative on the right hand side is provided in Appendix S1, while the third is simply $\phi$. The second partial derivative is obtained as

$$\begin{aligned}
\frac{\partial p_i}{\partial \phi} &= -\frac{\partial}{\partial \phi}\left\{\frac{1}{1+\phi\exp(\eta_i)}\right\}^{\phi^{-1}} \\
&= \left\{\frac{1}{1+\phi\exp(\eta_i)}\right\}^{\phi^{-1}} \\
&\quad \times \left[\log\left\{\frac{1}{1+\phi\exp(\eta_i)}^{\phi^{-1}}\right\}\frac{1}{\phi} + \frac{1}{\phi}\{1+\phi\exp(\eta_i)\}\{1+\phi\exp(\eta_i)\}^{-2}\exp(\eta_i)\right] \\
&= \left\{\frac{1}{1+\phi\exp(\eta_i)}\right\}^{\phi^{-1}}\frac{1}{\phi}\left[\frac{1}{\phi}\log\left\{\frac{1}{1+\phi\exp(\eta_i)}\right\} + \frac{1}{\exp(-\eta_i)+\phi}\right].
\end{aligned}$$

It follows that

$$\frac{\partial \ell_{\text{Bern}}^{\text{gcloglog}}}{\partial \log\phi} = \sum_{i=1}^{N}\left(\frac{y_i}{p_i}-1\right)\left[\frac{1}{\exp(-\eta_i)+\phi} - \frac{1}{\phi}\log\{1+\phi\exp(\eta_i)\}\right].$$

Moreover, the negative partial second derivative for $\beta$ is given by

$$
\begin{aligned}
-\mathcal{I}_{\log\phi,\beta} &= \sum_{i=1}^{N} \frac{\partial}{\partial \eta_i}\left(\frac{y_i}{p_i}-1\right)\left[\frac{1}{\exp(-\eta_i)+\phi}-\frac{1}{\phi}\log\left\{1+\phi\exp(\eta_i)\right\}\right]\frac{\partial \eta_i}{\partial \beta} \\
&= -\sum_{i=1}^{N} \frac{y_i}{p_i^2}\exp(\eta_i)\frac{1-p_i}{1+\phi\exp(\eta_i)}\left\{\frac{1}{\exp(-\eta_i)+\phi}+\log(1-p_i)\right\}\boldsymbol{x}_i \\
&+ \left(\frac{y_i}{p_i}-1\right)\Big[\{\exp(-\eta_i)+\phi\}^{-2}\exp\{-\eta_i\}-\{1+\phi\exp(\eta_i)\}^{-1}\exp\{\eta_i\}\Big]\boldsymbol{x}_i.
\end{aligned}
$$

Similarly, the negative second partial derivative for $\log\phi$ is given by

$$
\begin{aligned}
-\mathcal{I}_{\log\phi} &= \sum_{i=1}^{N} \frac{\partial}{\partial \phi}\left(\frac{y_i}{p_i}-1\right)\left[\frac{1}{\exp(-\eta_i)+\phi}-\frac{1}{\phi}\log\left\{1+\phi\exp(\eta_i)\right\}\right]\frac{\partial \phi}{\log \phi} \\
&= -\sum_{i=1}^{N} \frac{y_i}{p_i^2}(1-p_i)\left\{\log(1-p_i)+\frac{1}{\exp(-\eta_i)+\phi}\right\}^2 \\
&+ \left(\frac{y_i}{p_i}-1\right)[-\phi\{\exp(-\eta_i)+\phi\}^{-2}-\log(1-p_i)-\{1+\phi\exp(\eta_i)\}^{-1}\exp\{\eta_i\}].
\end{aligned}
$$

Letting $\boldsymbol{\Sigma}_\beta$ denote the (naive) asymptotic covariance matrix of a fitted GLM with fixed $\phi$, then

$$
\mathbb{V}_{\text{Bern}}^{\text{gcloglog}}(\hat{\boldsymbol{\beta}}) = \boldsymbol{\Sigma}_\beta + \frac{\boldsymbol{\Sigma}_\beta \mathcal{I}_{\beta,\log\phi}\mathcal{I}_{\beta,\log\phi}^{\top}\boldsymbol{\Sigma}_\beta}{\mathcal{I}_{\log\phi,\log\phi}-\mathcal{I}_{\beta,\log\phi}^{\top}\boldsymbol{\Sigma}_\beta\mathcal{I}_{\beta,\log\phi}}.
$$

In essence, since $\phi$ is a scalar, then we can leverage the Sherman-Morrison formula for a rank-one update to the naive asymptotic covariance matrix. The update additionally serves to reiterate that standard errors due to $\boldsymbol{\Sigma}_\beta$ alone are an underestimate of the standard error due to the joint information, given that the rank one update is positive when the joint information is positive definite. Finally, note that the expected information is straightforwardly derived from the two equations above, as it only requires minor simplification due to the (trivial) expectation $\mathbb{E}(y_i) = p_i$.

For models fitted with $\phi = 0$, we report the naive standard error, as the joint information is not positive definite in such cases.

Finally, note that the above derivation does not straightforwardly apply to more complex settings such as gcloglog link binomial GLMMs as considered in the case studies in Section 5. Instead, standard errors can be derived using the asymptotic covariance matrix provided by the `lme4` `R`-package, say, but using a finite differences approximation to the second partial derivative and second derivative for $\phi$

computed via the `numDeriv` R-package (Gilbert and Varadhan, 2019).

### S3 Additional result for the case studies

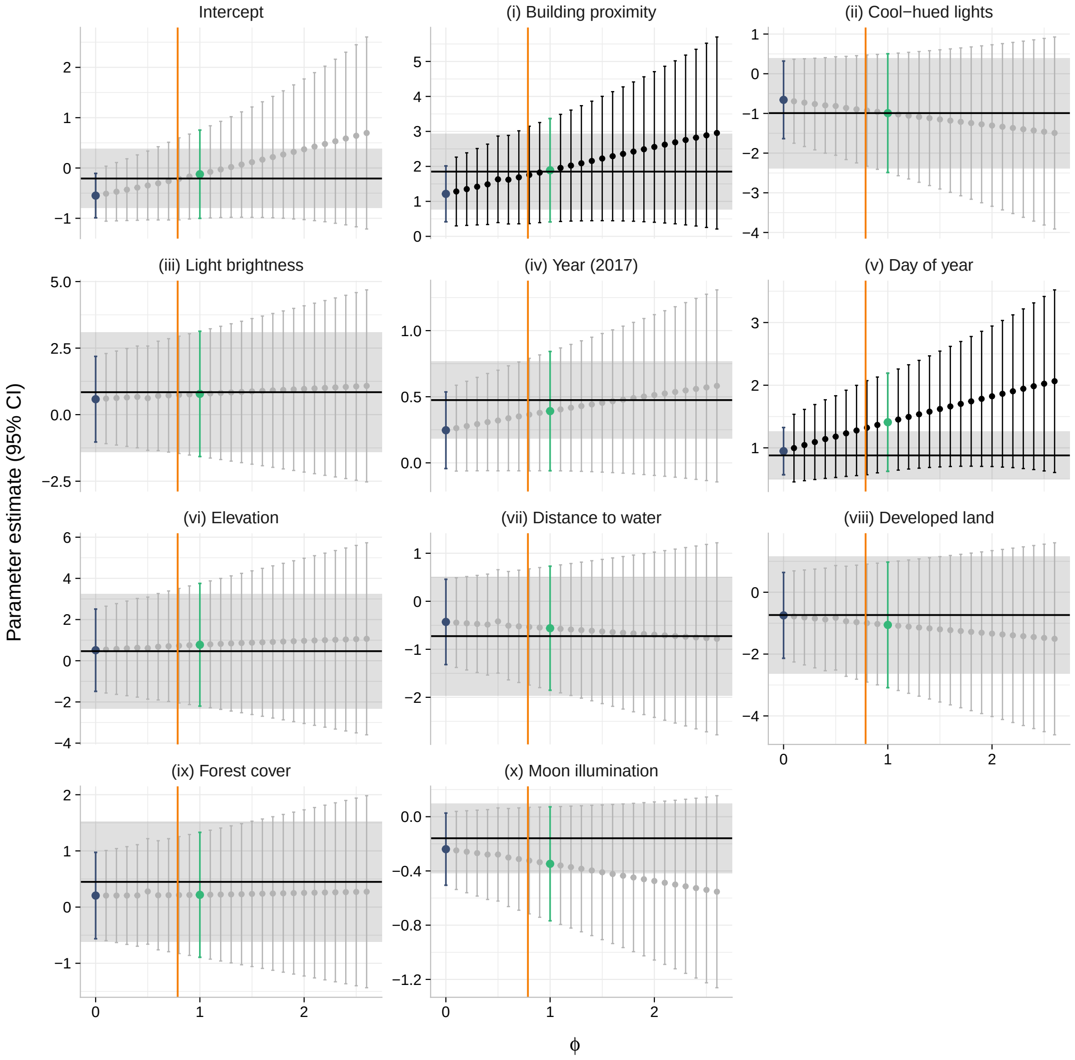


Figure S1: Sensitivity analysis for the gcloglog link GLMM for the application to the Bat data, based on varying the shape parameter $\phi$ within the range of its corresponding 95% confidence interval $(0, 2.642)$. The point estimates for the intercept along with the ten fixed effect covariates at each fitted value of $\phi$ are shown, with the bars showing the corresponding 95% confidence intervals. Confidence intervals that include zero, i.e., effects that are not statistically significant, are shown in grey. The solid horizontal line and shaded band in each panel mark the negative binomial GLMM estimate of $\phi$ and its corresponding 95% confidence interval for that estimate, while the cases of $\phi = 1$ (logit; green) and $\phi = 0$ (cloglog; blue) are also shown. Finally, the orange vertical line marks the profiled maximum likelihood estimate, $\hat{\phi} = 0.787$, for the gcloglog link GLMM.

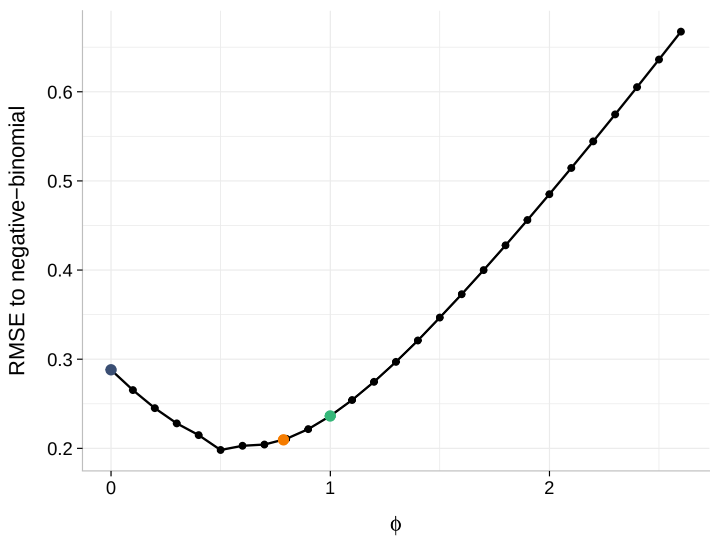


Figure S2: RMSE of the fixed parameter estimates between a gcloglog link binomial GLMM and the benchmark negative binomial GLMM for the application to the Bat data, varying as a function of $\phi \in \{0, 0.1, \ldots, 2\}$. The points correspond to $\phi = 0$ (blue; cloglog), $\phi = 1$ (green; logit) and $\hat{\phi} = 0.787$ (orange; gcloglog).

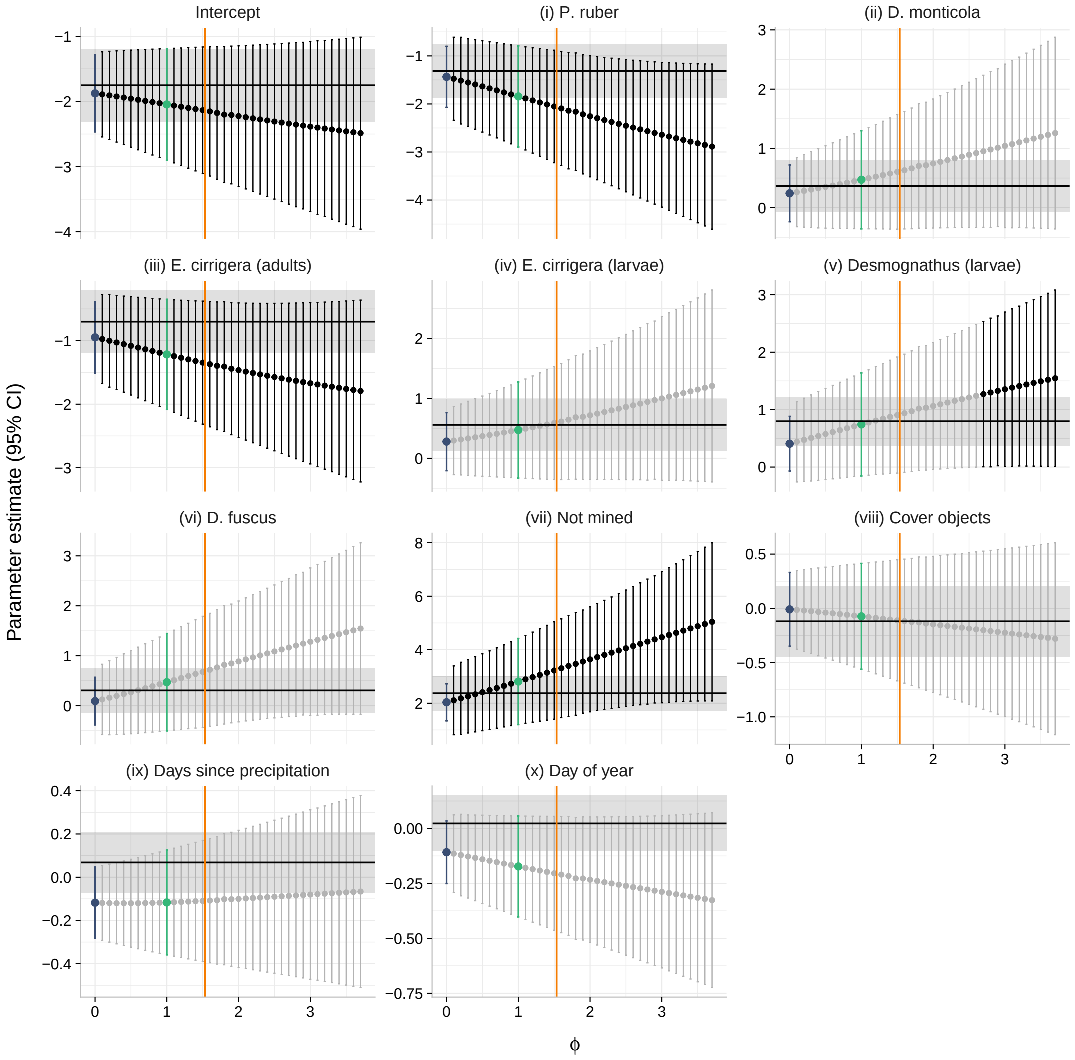


Figure S3: Sensitivity analysis for the gcloglog link GLMM in the application to the Salamandar data, based on varying the shape parameter $\phi$ within the range of its corresponding 95% confidence interval ranging $(0, 3.787)$. The point estimates for the intercept along with the ten fixed effect covariates at each fitted value of $\phi$ are shown, with the bars showing the corresponding 95% confidence intervals. Confidence intervals that include zero, i.e., effects that are not statistically significant, are shown in grey. The solid horizontal line and shaded band in each panel mark the negative binomial GLMM estimate of $\phi$ and its corresponding 95% confidence interval for that estimate, while the cases of $\phi = 1$ (logit; green) and $\phi = 0$ (cloglog; blue) are also shown. Finally, the dotted vertical line marks the profiled maximum likelihood estimate, $\hat{\phi} = 1.534$, for the gcloglog link GLMM.

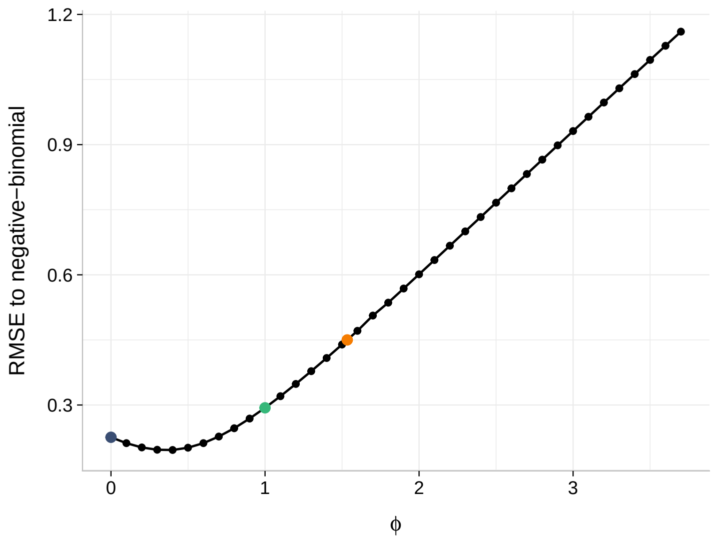


Figure S4: RMSE of the fixed parameter estimates between a gcloglog link binomial GLMM and the benchmark negative binomial GLMM for the application to the Salamandar data, varying as a function of $\phi \in \{0, 0.1, \ldots, 2\}$. The points correspond to $\phi = 0$ (blue; cloglog), $\phi = 1$ (green; logit) and $\hat{\phi} = 1.534$ (orange; gcloglog).
